# Protocol for muscle fiber type and cross-sectional area analysis in cryosections of whole lower mouse hindlimbs

**DOI:** 10.1101/2024.08.22.608909

**Authors:** Edmund Battey, Dipsikha Biswas, Mathieu Dos Santos, Pascal Maire, Kei Sakamoto

## Abstract

We outline a robust, simple, and cost-effective method to simultaneously visualize all mouse lower hindlimb skeletal muscles. We describe procedures for orientating the whole lower hindlimb in gum tragacanth prior to freezing, simplifying the proceeding experimental steps, and enhancing the clarity and comprehensiveness of characterizations. We then detail steps for quantifying muscle fiber size and fiber type characteristics in a single cryosection using immunofluorescence and histochemistry. This protocol can be applicable for commonly used histological and (immuno) histochemical evaluations such muscle degeneration/regeneration, fibrosis, immune cell infiltration, enzymatic activity and glycogen content.

**Highlights:**

- Embedding and orientating mouse lower hindlimb in tragacanth before freezing enhances clarity and simplifies steps.
- Simultaneously visualizing myofibers in different muscle compartments enables efficient and comprehensive evaluations of morphology and physiological characteristics.
- Single cryosection analysis of myofiber size and types with immunofluorescence.
- Applicable for evaluating muscle degeneration, fibrosis, and immune infiltration.

**Graphical abstract:** 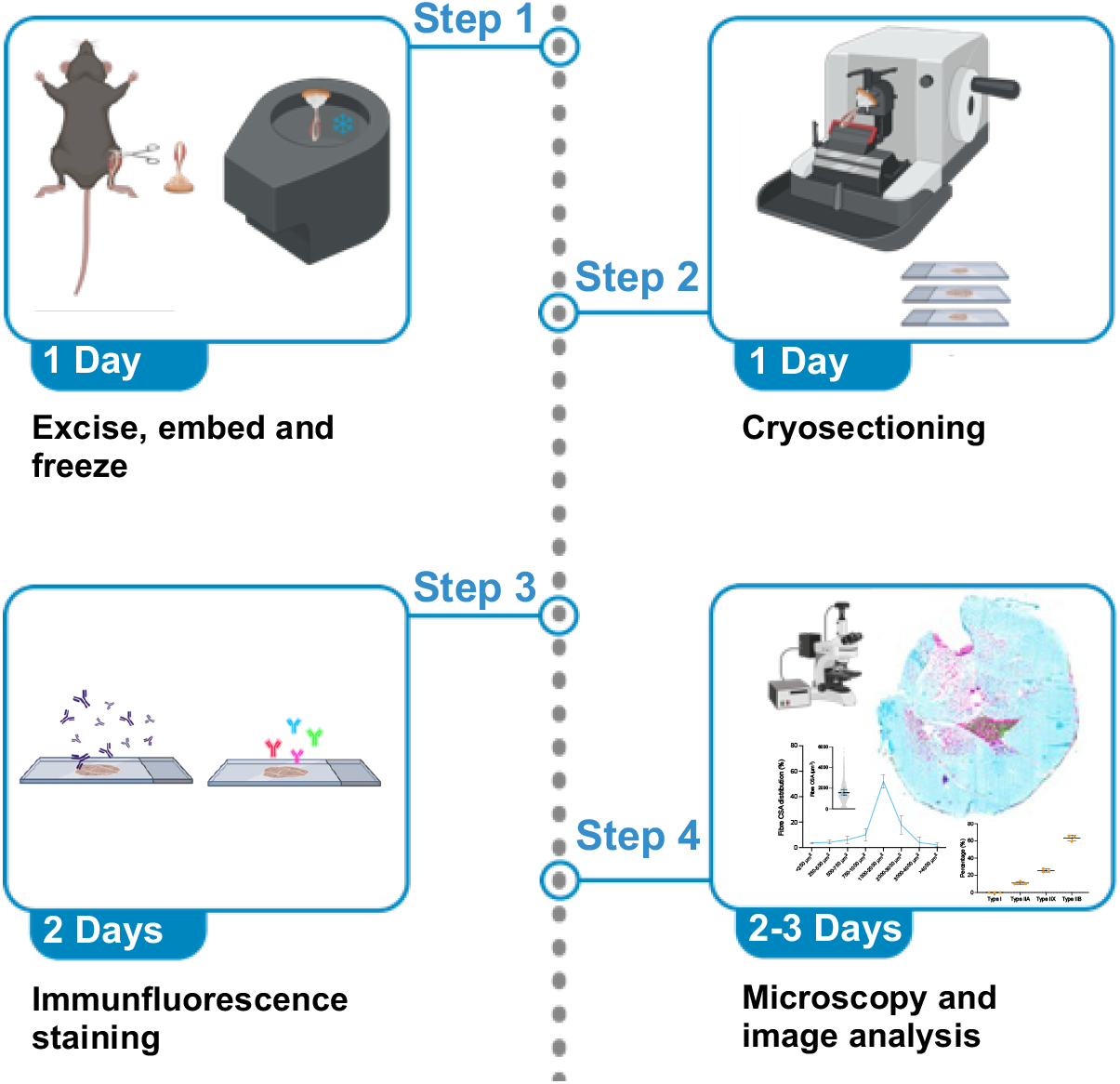

## Before you begin

Skeletal muscle is a complex tissue that plays a critical role in locomotion, posture, and metabolism. It is composed of different fiber types, each characterized by distinct metabolic profiles and contractile properties due to the expression of specific sarcomeric proteins among which myosin heavy chain (MyHC) isoforms^1^. These fiber types, classified as type I, IIa, IIx, and IIb, exhibit varying rates of contraction, fatigue resistance, and oxidative capacity, making them essential for the diverse functional requirements of skeletal muscle^1^.

Accurate identification and quantification of muscle fibre types are fundamental for understanding muscle function, adaptation, and response to various stimuli, such as exercise, aging, and disease^1–6^. Moreover, studying fiber type distribution in muscle-related pathologies can provide valuable insights into exercise performance, insulin sensitivity, disease progression and therapeutic interventions^7,8^.

Traditional methods for analyzing muscle fibre types involve time-consuming procedures, often necessitating multiple cryosections from different muscles ^9^. Therefore, a method that allows for the simultaneous investigation of all lower hindlimb muscles in a single cryosection would save time and resources and ensure concurrent analysis of different muscles in a single sample.

In this study, we introduce a robust and simple approach that addresses these challenges by enabling efficient and comprehensive analysis of muscle fibre types and sizes in mouse lower hindlimb cryosections. Our method leverages immunofluorescence staining, which permits the visualization of multiple fiber types simultaneously. By optimizing the embedding, freezing, and immunofluorescence staining protocol, we achieve reliable and specific labelling of each fiber type in the entire leg musculature.

The protocol below describes the specific steps for skeletal muscle fiber typing and cross-sectional area analysis. Additionally, this protocol can also be applied for other commonly used histological and (immuno) histochemical evaluations such muscle degeneration/regeneration, fibrosis, immune cell infiltration and glycogen content.

## Institutional permissions

Animal experiments were conducted in accordance with the European directive 2010/63/EU of the European Parliament and of the Council of the protection of animals used for scientific purposes. Ethical approval was given by the Danish Animal Experiments Inspectorate (license number #2021-15-0201-00884). Wild type C57BL/6NTac male mice were obtained from Taconic Biosciences and housed in the animal facility at the Faculty of Health and Medical Sciences (University of Copenhagen). All the animals were kept and maintained according to local regulations under a light/dark cycle of 12 h and had free access to a standard chow diet.

## Key resources table

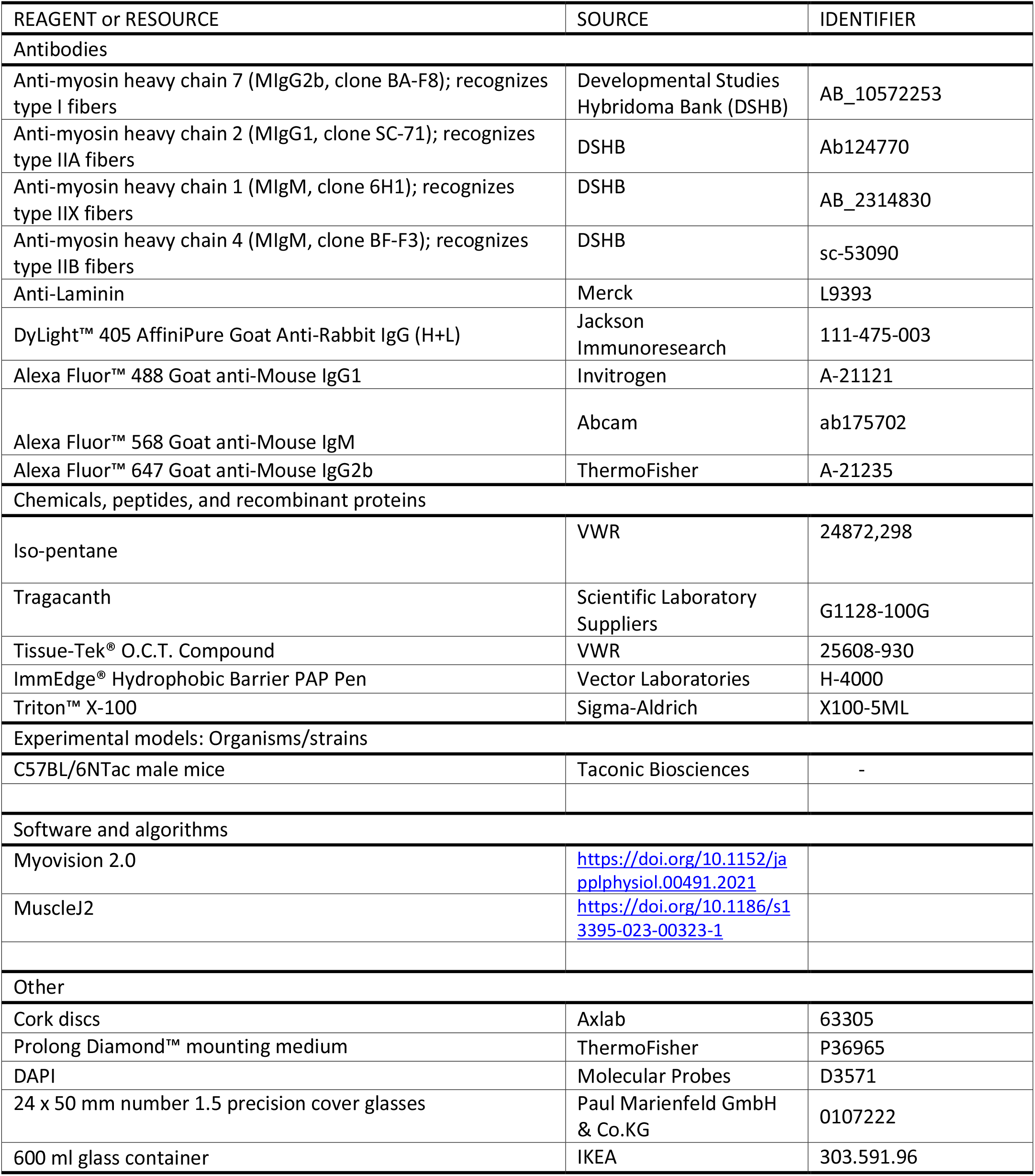

## Materials and equipment

## Tragacanth solution for embedding

- Make up 6 % tragacanth solution in double distilled water (ddH_2_O ) or Milli-Q water or embedding whole lower hindlimb prior to freezing. 50 – 100 ml final volume will be sufficient for 5-10 mice.
- Dissolve 6 g tragacanth for every 100 ml ddH_2_O by microwaving in short bursts and stirring intermittently.
- Store tragacanth in 50 ml falcon tubes on ice or at 4°C for up to 2 weeks until the embedding step.

## Antibody solutions

***Note:****antibody concentrations can be optimized depending on titer/batches*

**Primary antibody cocktail 1 (diluted in PBS)**

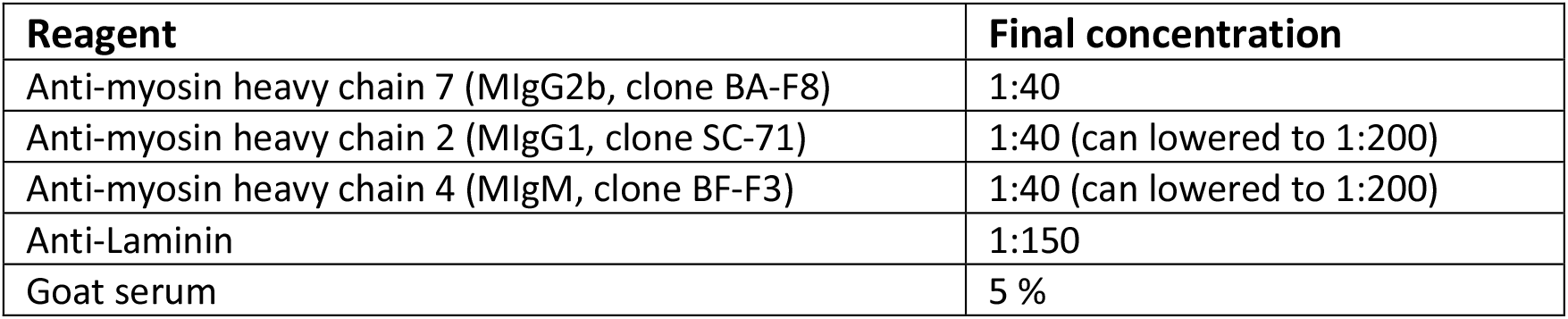

| Reagent | Final concentration |
| --- | --- |
| Anti-myosin heavy chain 7 (MlgG2b, clone BA-F8) | 1:40 |
| Anti-myosin heavy chain 2 (MlgG1, clone SC-71) | 1:40 (can lowered to 1:200) |
| Anti-myosin heavy chain 4 (MlgM, clone BF-F3) | 1:40 (can lowered to 1:200) |
| Anti-Laminin | 1:150 |
| Goat serum | 5 % |

**Primary antibody cocktail 2 (diluted in PBS)**

| Reagent | Final concentration |
| --- | --- |
| Anti-myosin heavy chain 1 (MlgM, clone 6H1) | 1:40 |
| Anti-Laminin | 1:150 |
| Goat serum | 5 % |

**Secondary antibody cocktail 1 (diluted in PBS)**

| Reagent | Final concentration |
| --- | --- |
| DyLight™ 405 AffiniPure Goat Anti-Rabbit IgG (H+L) | 1:800 |
| Alexa Fluor™ 488 Goat anti-Mouse IgG1 | 1:800 |
| Alexa Fluor™ 568 Goat anti-Mouse IgM | 1:800 |
| Alexa Fluor™ 647 Goat anti-Mouse IgG2b | 1:800 |
| Goat serum | 5 % |

**Secondary antibody cocktail 2 (diluted in PBS)**

| Reagent | Final concentration |
| --- | --- |
| DAPI | 1:800 |
| Alexa Fluor™ 488 Goat anti-Rabbit IgG (H+L) Cross-Adsorbed Secondary Antibody | 1:800 |
| Alexa Fluor™ 568 Goat anti-Mouse IgM | 1:800 |
| Goat serum | 5 % |

## CRITICAL

Isopentane is harmful and toxic to breathe in – use under a fume hood. Use protective gloves, eye protection and keep away from heat/ignition sources.

## CRITICAL

Direct contact with liquid nitrogen can cause severe harm, including cryogenic burns, frostbite, and eye injuries. Wet skin is especially susceptible to freezing. Use protective gloves and eye protection.

## Step-by-step method details

## Pre-cool isopentane for freezing

Timing: 15 min

1. Half-fill a glass or metal container with isopentane (approximately 500 ml capacity)
2. Fill a larger polystyrene container with a volume of liquid nitrogen that will reach approximately half-way up the container with isopentane
3. Place the container with isopentane into the larger container with liquid nitrogen and allow to cool for approximately 2 minutes until the vapour starts to disappear and a white precipitate forms on the bottom (Figure 2A).

**Figure 1.**
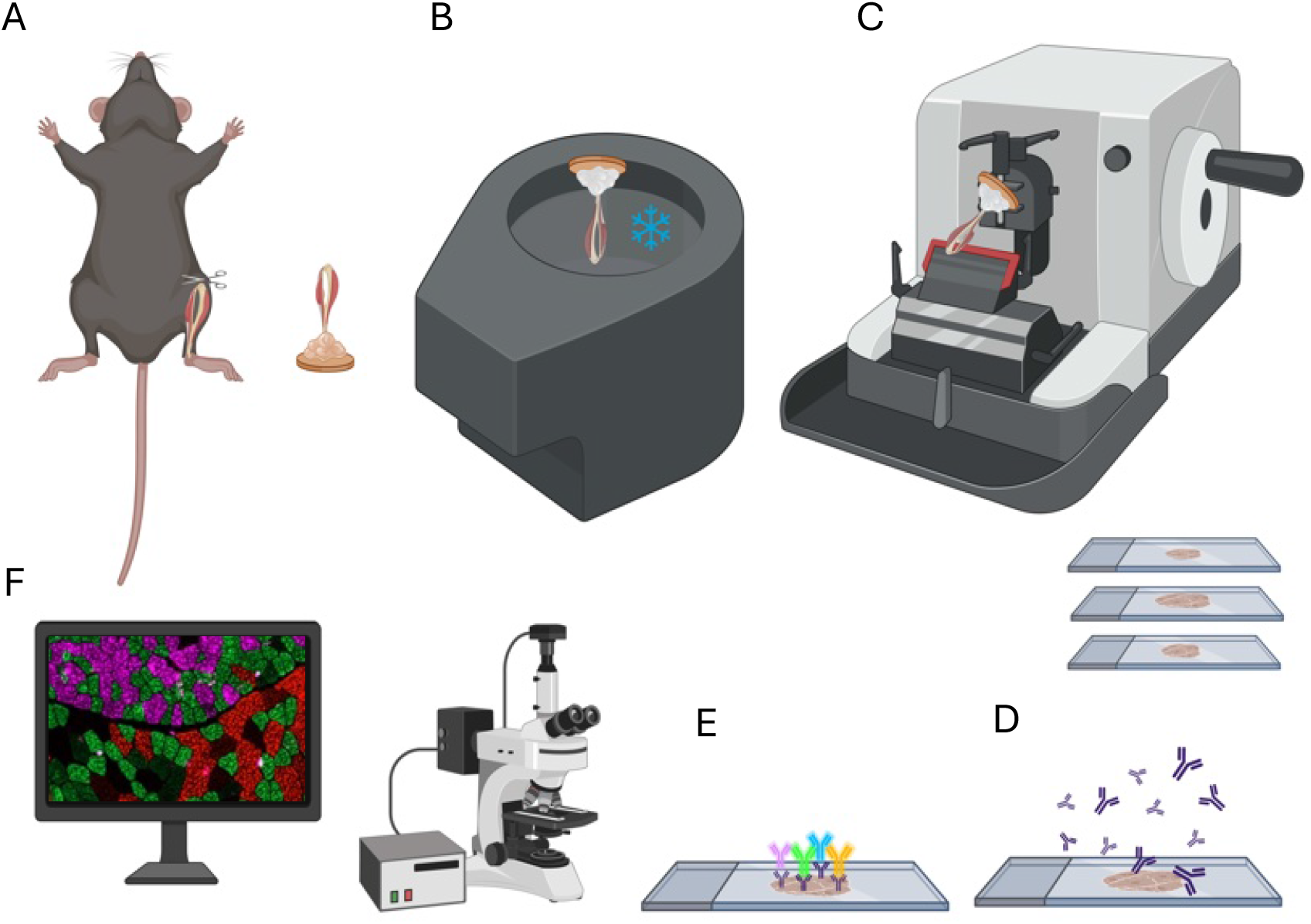
Overview of protocol steps including sample excision, freezing, cryosectioning, immunohistochemistry and microscopy. (A) Excise lower hindlimb and embed in tragacanth. (B) Freeze in liquid nitrogen-cooled isopentane. (C-E) Cryosection and apply primary antibodies directed against Myosin Heavy Chain (MHC) isoforms for fiber typing and Laminin to mark fiber borders. Use secondary antibodies conjugated to fluorophores for visualization of proteins. (F) Image sections using a widefield or confocal fluorescence microscope.

**Figure 2.**
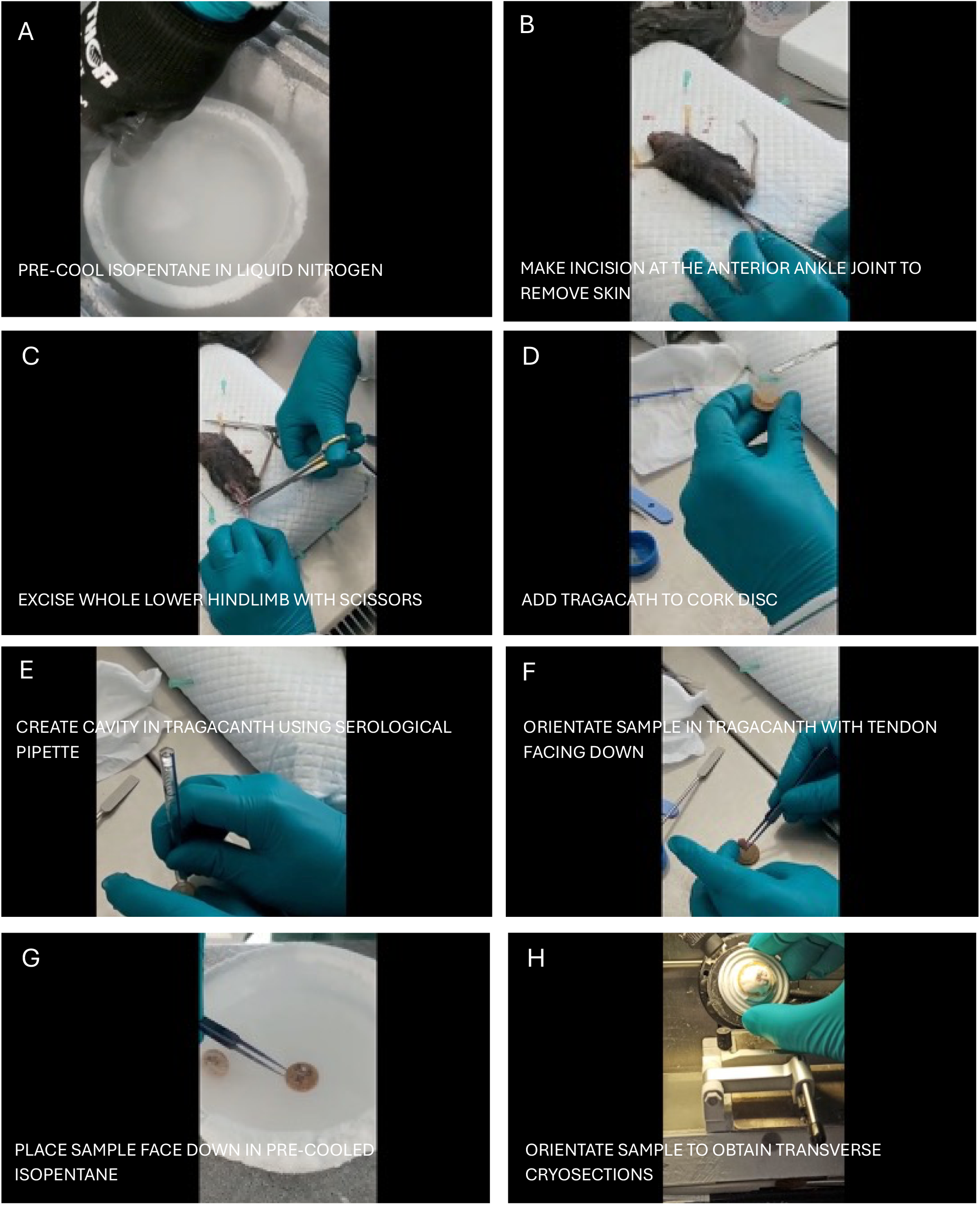
Step-by-step protocol for excising, embedding, freezing, and cryosectioning mouse whole lower hindlimb. (A) Pre-cooled isopentane in liquid nitrogen. (B) Make an incision at the anterior ankle joint to remove the skin. (C) Excise the whole lower hindlimb with scissors. (D) Add tragacanth to cork disc. (E) Create a cavity in tragacanth using a serological pipette. (F) Orientate sample in tragacanth with tendon facing down. (G) Place sample face down in pre-cooled isopentane. (H) Orientate sample to obtain transverse cryosections.

## CRITICAL

Ensure that the isopentane does not over-solidify prior to freezing the muscle. Periodic removal of the isopentane from the liquid nitrogen may be necessary.

## Excision of lower hindlimb

Timing: 3 min per leg

Excision

This step results in obtaining the mouse lower hindlimb ready for freezing.

4. Following the protocol outlined by Moyle and Zammit (2014)^10^ position the mouse to maximize ease and efficiency of excision (Figure 2B).
  i. Place forelimb paws above the head of the mouse and pin to a polystyrene board.
  ii. Pin the dorsal paw of the left hindlimb parallel to the midline and pin the contralateral hindlimb at a 45-degree angle away from the mid-line.
  iii. Pull the tail under the left hindlimb and pin to the board to maximize stability and access to the anterior hindlimb muscles.
5. Remove the skin covering the leg of the mouse.
  i. To minimize contamination of samples with hair, spray the left hindlimb with 70% ethanol.x
  ii. Make an incision using scissors into the skin at the anterior side of the ankle joint (Figure 2B).
  iii. Cut the skin up to the hip and pull away to expose the hindlimb musculature.
6. Use large, sharp scissors to cleanly cut the leg below the patella (Figure 2C).
7. Cut the hindlimb paw of the mouse, leaving some of the distal tendons attached to the leg to help maintain the orientation and arrangement of the muscles.

## Optional

It is possible to use the forelimb or the proximal part of the hindlimb to perform cryosections^11^.

## Embedding and freezing lower hindlimb

Timing: 5 min per leg

This step accomplishes orientation of the sample in tragacanth and freezing in liquid nitrogen-cooled isopentane, thereby preparing the sample for cryosectioning.

8. Apply gum tragacanth to cork disc, or alternatively optimal cutting temperature (OCT) compound in a cryomold, and create a cavity (Figure 2D-E).
  i. Use a spatula to place approximately 2 x 2 cm of tragacanth on the cork disc, or if using a cryomold, fill the mold with approximately 2 cm layer of OCT
  ii. Use a serological pipette to create a cavity approximately 1 cm in diameter (**Troubleshooting 1**).
9. Place the hindlimb tendon-down into the cavity (Figure 2F).
  i. Ensure that the bottom of the hindlimb is covered in tragacanth or OCT to avoid it being displaced during freezing in isopentane.
  ii. Orientate the hindlimb so that the angle is perpendicular to the cork disc or cryomold.
10. Freeze the embedded hindlimb by placing it face-down in isopentane for 1-2 min (Figure 2G).
11. Transfer embedded hindlimb to dry ice until isopentane evaporates (Figure 2H).

## Optional

Store samples in small ziplock bags in the -80°C freezer until cryosectioning is to be completed.

## Cryosectioning

12. Set the cryostat chamber temperature to -20°C and object temperature to -18°C.
13. Place a new blade in the cryostat blade holder for clean and precise sectioning.
14. Take embedded samples from the -80°C freezer and transport them on dry ice to the cryostat.
15. Place the samples in the cryostat chamber, close the lid, and let it equilibrate to the cryostat temperature for 20-30 minutes.
16. Use OCT solution to stick the cork disc sample to the sample chuck.
17. Orientate the sample in the cryostat sample so that it perpendicular to the blade.

## CRITICAL

the muscle fibers must be orientated perpendicular to the cryostat blade to ensure reliable quantification of muscle fiber parameters (especially cross-sectional area). At this stage, sections can be checked for correct orientation using a stereoscope. Fiber borders should appear circular, rather than elongated (see Figure 4, for example).

18. Cut the sample until it flattens and reaches the section of the leg with major muscles.
19. Set the section thickness on the cryostat to 12 μm and collect serial sections on Superfrost™ Plus Microscope Slides. A 12 μm section thickness balances detail and stain penetration, and minimizes artifacts created by the cutting process, which can occur if the sections are too thin.
20. Let the collected sections dry out at room temperature for at least 30 min.
21. Place them in a slide box, place the slide box in a bag to avoid moisture accumulation, and store at -80°C until immunohistochemistry steps.

## Optional

To enhance the condition of sections for improved clarity, the tissue can be trimmed with an old razor blade and replaced with a new one when the section needs to be collected into slides.

## Immunohistochemistry

22. Thaw slides at room temperature, air dry for 15 min, draw a hydrophobic ring around sections using a hydrophobic barrier pen (for example, ImmEdge® Hydrophobic Barrier PAP Pen).
23. Rehydrate in PBS for 10 min either by submersion in a Coplin jar or by using pipette to add enough PBS to cover the section.

## Optional

add a permeabilization step by dipping the slide very quickly in 0.3% Triton™ X-100/PBS and removing it immediately. This allows the primary antibodies to be distributed better on the sections.

24. Block sections in 5% goat serum in PBS for 10 min to reduce non-specific binding of antibodies. Use a pipette to add approximately 200 μl of the blocking solution onto the sections.
25. To enable comprehensive analysis of muscle fiber types across various muscles simultaneously, stain sections using specific antibodies directed against MYH7, MYH2, and MYH4 isoforms (Figure 2; Primary antibody cocktail 1). These antibodies are used because they are each different mouse isotypes and can therefore be detected simultaneously by isotype-specific secondary antibodies. This staining strategy facilitates the distinct visualization of type I, type IIA, and type IIB fibers, while the absence of staining indicates type IIX fibers. Primary antibody incubation should be carried out overnight at 4°C.

## Optional

on a separate slide, use antibodies directed against MYH1 and Laminin (Primary antibody cocktail 2).

26. Wash sections with PBS (3 x 5 min) and incubate with secondary antibodies on a shaker set to 200 rpm (Secondary antibody cocktail 1 or Secondary antibody cocktail 2) for 1 hour.
27. Wash sections with PBS (3 x 5 min) and mount slides in the anti-fade reagent, Prolong Diamond™ and leave to cure at room temperature for >24 hours before imaging.

## Imaging and analysis

28. Acquire tiled images of lower hindlimb sections using a widefield or confocal microscope equipped with a 10× air objective and imaging software such as Zen Pro.
29. Calculate fiber type proportions and fiber cross-sectional area manually or semi-automatically using plugins/ software such as MuscleJ2^12^ or Myovision 2.0^13^. If quantifying fiber types manually, use the multi-point tool in FIJI to record each the counts of each fiber type. Divide the number of each fiber types by the total number of fibers, and multiply by 100 to obtain fiber type proportions as percentages.

## Expected outcomes and quantifications

This method permits visualization and quantification of fiber types (Figure 3) and cross-sectional area (CSA) (Figure 4) in the same section and has previously been applied on a transgenic mouse model developed by our group^14^. Fiber typing of extensor digitorum longus (EDL) and soleus can be carried out manually using FIJI^15^ (Figure 3C, 3E) or semi-automatically using purpose-built software such as MuscleJ2^12^ or Myovision 2.0^13^. Here, fiber typing was completed manually using FIJI and Myovision 2.0^13^ was used to semi-automatically quantify muscle fiber CSA of EDL and soleus (Figure 4C, 4E).

**Figure 3.**
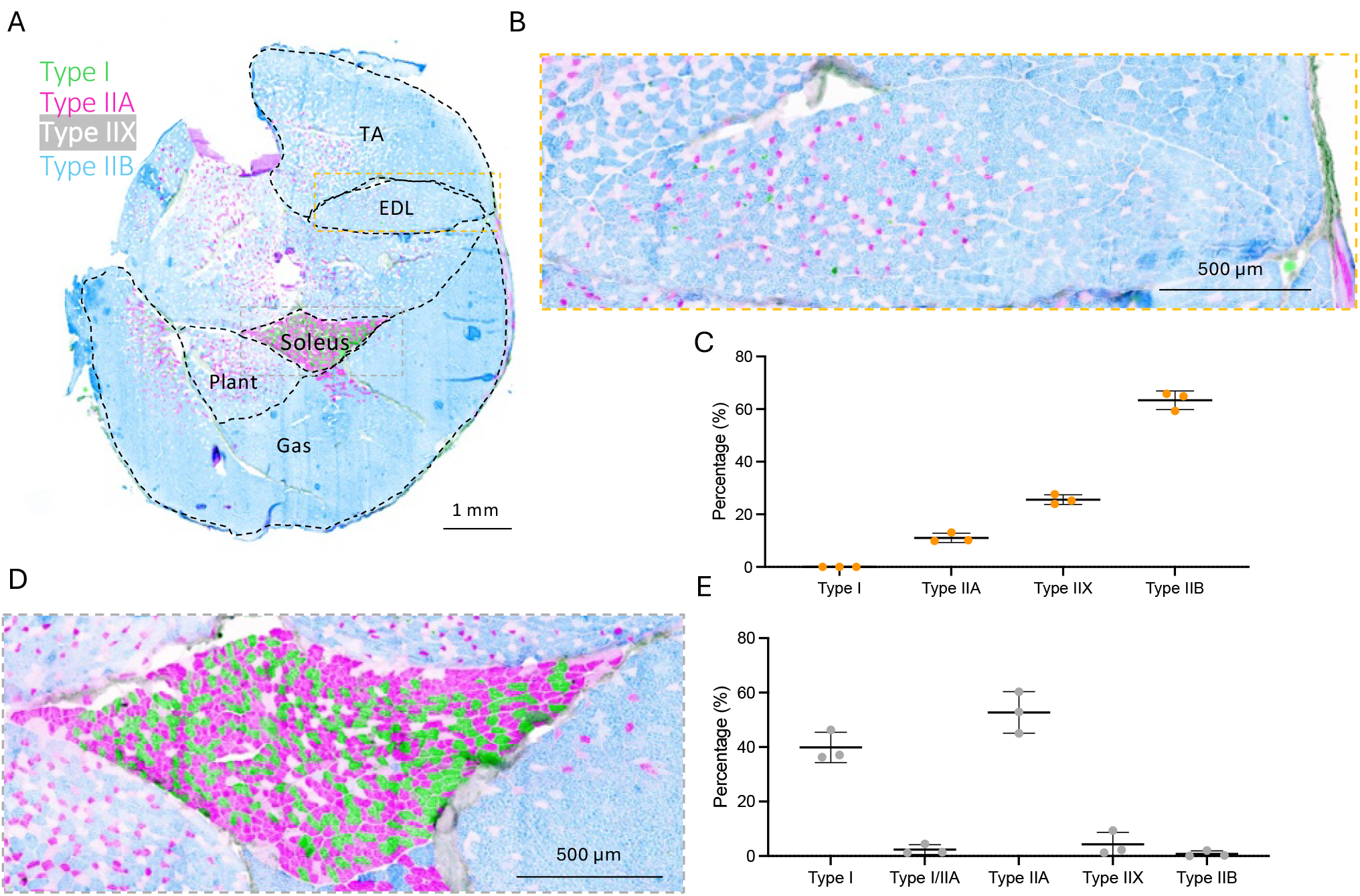
Representative image and quantifications of transverse cryosection mouse lower hindlimb stained with antibodies to visualize fiber types. A) Transverse cross-section of the lower hindlimb showing different muscle groups: tibialis anterior (TA), extensor digitorum longus (EDL), soleus, plantaris (Plant), and gastrocnemius (Gas). Muscle fibers are color-coded for different fiber types: Type I (green), Type IIA (pink), Type IIX (white), and Type IIB (blue). (B) Magnified view of the EDL muscle showing the distribution of different fiber types. (C) Quantification of fiber type distribution in EDL muscle. (D) Magnified view of the soleus muscle with fiber type distribution. (E) Quantification of fiber type distribution in the soleus muscle. (F) Quantification of fiber type distribution in the EDL muscle.

**Figure 4.**
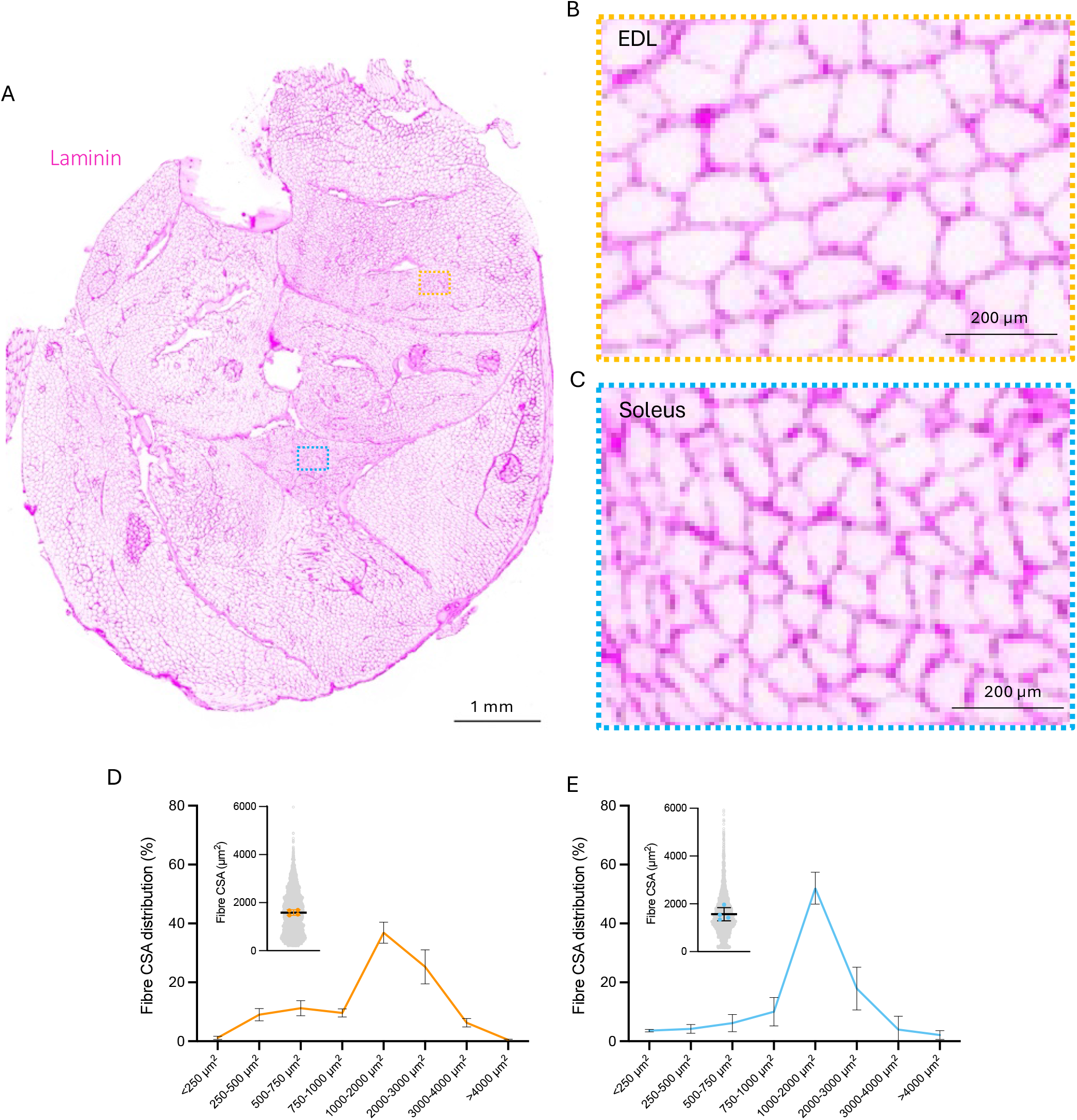
Representative images and quantifications of mouse lower hindlimb transverse cryosection stained with Laminin antibody to visualize fiber borders for cross-sectional area analysis. (A) Overview of the lower hindlimb cross-section stained with anti-Laminin antibody to visualize fiber borders (magenta), with Extensor Digitorum Longus (EDL) and soleus regions of interest marked by orange and blue boxes, respectively. (B-C) Magnified view of the EDL and soleus areas marked by the orange and blue boxes, showing detailed fiber borders. (D) Corresponding cross-sectional area quantification for the orange box region. (E) Magnified view of the area marked by the blue box, showing detailed fiber borders. (E) Corresponding cross-sectional area quantification for the blue box region.

Quantitative assessment of fibre type proportions within the EDL muscle were 11.05 ± 1.8 % Type IIA, 25.6 ± 1.9 % Type IIX, and 63.4 ± 3.5 % Type IIB fibres (Figure 3B-C). In the soleus muscle, fibre types were as follows: 39.9 ± 5.6 % Type I, 2.4 ± 1.8 % Type I/IIA hybrid, 52.7 ± 7.6 % Type IIA, 4.3 ± 4.4 % Type IIX, and 0.7 ± 1.1 % Type IIB fibres (Figure 3D-E). Fiber CSA of soleus and EDL were similar, as previously reported in mice of a similar age^16^, with EDL having a greater proportion of larger fibers compared to soleus (Figure 4). These fiber CSA and fiber typing data are comparable with previous reports^16–20^.

## Limitations

While the proposed method offers substantial benefits there are potential limitations and room for improvement. Achieving uniform staining intensity across different muscle types might present a challenge due to variations in tissue properties and fiber densities. Therefore, optimizing staining conditions, including antibody concentrations and incubation times may be necessary. Improvements incorporating automated image acquisition and advanced image analysis techniques could enhance the speed and accuracy of fiber type quantification. Additionally, integrating molecular markers with fiber type staining could provide insights into gene expression patterns associated with specific muscle fiber types, further enriching the analysis. Finally, this method uses all muscles of the lower hindlimb, meaning the muscles cannot be used for other analyses. However, the contralateral leg can be used for such analyses.

## Troubleshooting

## Problem 1: unable to create cavity in tragacanth

Pressing serological pipette into tragacanth does not result in creation of cavity

## Potential solution

Avoid warming tragacanth and keep on ice until it is ready to be used.

## Problem 2: isopentane solidifies

Isopentane cools too much and solidifies, preventing homogenous and quick freezing.

## Potential solution

Temporarily remove the container with isopentane from the liquid nitrogen until it becomes liquid, then return to liquid nitrogen until white precipitates form at the bottom of the container.

## Resource availability

## Lead contact

Edmund Battey

## Materials availability

There were no newly generated materials for use in this protocol.

## Data and code availability

Data from the authors is available upon request. No code generated for the protocol.

## Acknowledgments

The Novo Nordisk Foundation Center for Basic Metabolic Research is an independent Research Center based at the University of Copenhagen, Denmark, and partially funded by an un-conditional donation from the Novo Nordisk Foundation (Grant number: NNF23SA0084103). M.D.S. is supported by a K99 grant from the National Institutes of Health (NIAMS, K99AR084549).

## Author contributions

Edmund Battey carried out experiments, data analysis, data visualization and wrote the manuscript. Dipsikha Biswas carried out experiments and edited the manuscript. Mathieu Dos Santos contributed to protocol development, experimental design and edited the manuscript. Pascal Maire and Kei Sakamoto supervised the work and edited the manuscript.

## Declaration of interests

The authors declare no competing interests.

## Figure legends

**Methods Video S1: Step-by-step video of excising, embedding, freezing, and cryosectioning mouse whole lower hindlimb in tragacanth on cork discs**

**Methods Video S2: Step-by-step video of excising, embedding, freezing, and cryosectioning mouse whole lower hindlimb in OCT compound in cryomolds**

## Notes

### Competing Interest Statement

The authors have declared no competing interest.

